# Transoceanic pathogen circulation in the age of sail and steam

**DOI:** 10.1101/2024.01.01.573813

**Authors:** Elizabeth N. Blackmore, James O. Lloyd-Smith

## Abstract

In the centuries following Christopher Columbus’s 1492 voyage to the Americas, transoceanic travel opened unprecedented pathways for global pathogen circulation. Yet no biological transfer is a single, discrete event. We use mathematical modeling to quantify historical risk of shipborne pathogen introduction, exploring the respective contributions of journey time, ship size, population susceptibility, transmission intensity, density dependence, and pathogen biology. We contextualize our results using port arrivals data from San Francisco, 1850–1852, and from a selection of historically significant voyages, 1492–1918. We offer numerical estimates of introduction risk across historically-realistic ranges of journey time and ship population size, and show that both steam travel and shipping regimes that involved frequent, large-scale movement of people substantially increased risk of transoceanic pathogen circulation.

## Introduction

In the centuries following Christopher Columbus’s 1492 journey to the Americas, transoceanic voyages opened unprecedented pathways in global pathogen circulation. In 1962, historian Woodrow Borah described the changes that followed as near-immediate; in his account, previously isolated regions such as the Americas and the Pacific “received within a few decades the united impact of all the diseases that could be spread” [1]. This narrative of rapid—and inevitable— pathogen transfer continues to shape some popular histories of global infectious disease [2]. Yet while transoceanic shipping was indeed a pivotal ecological force, the transformations that followed took substantially longer than decades. Global pathogen transfer was—and is—a centuries-long process.

Sixty years on, scholars in both the humanities and the sciences have charted the slow globalization of infectious disease. In the 1970s, pioneering environmental histories such as Alfred Crosby’s “Columbian Exchange”, Emmanuel Le Roy Ladurie’s “microbial unification of the world”, and William McNeill’s “common market of microbes” expanded the scope of Borah’s analyses to show that first introductions of “Old World” pathogens into previously isolated regions spanned one or two centuries following European arrival [3–5]. Subsequent historians have shown that pathogen exchange across the Atlantic, Pacific, and Indian oceans occurred slowly, with some introductions causing only transient outbreaks [2,6–13]. These outcomes were highly contingent on local human processes such as trade, warfare, and colonialism. In parallel, disease ecologists have shown that acute pathogens such as measles and influenza require large human populations for endemic local establishment [14–17], and that in smaller populations, continued circulation depends on regular introductions from “source” populations [18,19]. These metapopulation dynamics were as critical to historical pathogen dynamics as they are today [20–25]. Throughout the eighteenth century, the city of Boston, Massachusetts experienced decades-long intervals between smallpox outbreaks [26,27], while the much larger city of London experienced smallpox cases every year since records began in 1664 [27]. Sporadic introductions were, and are, extremely impactful – particularly in populations with no prior immunity [28–30]. Sustained “microbial unification”— the transition from intermittent introduction to continuous global circulation—required regular human movement and, with it, continued introduction and re-introduction of pathogens.

This raises an ecological question. How easily did infectious diseases survive the weeks- or months-long voyages necessary for transoceanic pathogen transfer in the age of sail and steam? There is good reason to expect that transoceanic pathogen introduction under these conditions was far from assured – particularly for fast-burning respiratory infections such as smallpox, measles, and influenza. As late as the 1850s, a sail voyage from Liverpool to New York City could take 5-6 weeks [31], while journeys from the UK to Australia could take 3-4 months [32]. Between lengthy periods at sea, short infection generation times, and intense shipboard transmission, fast-burning “crowd diseases” could rapidly exhaust all susceptible people on board and go extinct long before a vessel reached port, leaving no pathogen to introduce.

We explore the mechanics of shipborne pathogen transfer using the toolkit of contemporary theoretical ecology. We present a stochastic SEIR model which quantifies the probability of an outbreak lasting a given duration in a closed population. We consider the relative contributions of a broad range of factors to outbreak duration, including pathogen natural history, transmission intensity and density dependence, population size, and population susceptibility. Finally, we use port arrivals data from Gold Rush-era San Francisco, California, 1850–1852, to explore the implications of variation in journey length, ship size, and natural history for pathogen circulation in the specific context of the Pacific. As part of this, we explore the impact of the advent of steam travel in the nineteenth century – a technological revolution that routinely cut journey times by a factor of two or more [31–33].

The idea that many shipboard outbreaks ended long before a vessel’s arrival is intuitive. These processes have been considered qualitatively, both by scientists [33,34] and by historians [3,10,13,35]. Paterson et al. [36] have also modeled the specific case of shipborne measles introductions to Australia during the nineteenth century. A more general quantitative analysis can offer sharper insight into the contours of global disease history, and can aid in building broader structural histories of infectious disease [9,13,37,38]. It can also reveal new patterns in seemingly disparate disease introductions. Our results indicate that shipborne pathogen introductions were neither trivial nor inevitable. Ships were not simple pathogen vectors: they were populations. The extinction and survival dynamics of pathogens on ships were complex population biological processes, contingent on pathogen natural history and host population size, composition, and mixing patterns. Thus, the history of transoceanic disease introduction is a story both of fundamental pathogen biology, and of human economics, technology, and behavior.

Theoretical modelling can reveal how these forces interacted to shape global disease transmission.

## Results

### Basic Dynamics

Transoceanic pathogen introduction requires a chain of infections that lasts at least as long as a ship’s journey time. To investigate the basic dynamics of shipboard outbreak duration, we simulate outbreaks in a fully susceptible population (*N* = 100) using a hypothetical pathogen which has characteristics typical of acute respiratory viruses (mean incubation and infectiousness periods of 5 days each) (Figure 1). We define outbreak duration as the time until nobody on board ship is infected with the pathogen, that is, until *E* = *I* = 0.

**Fig. 1.**
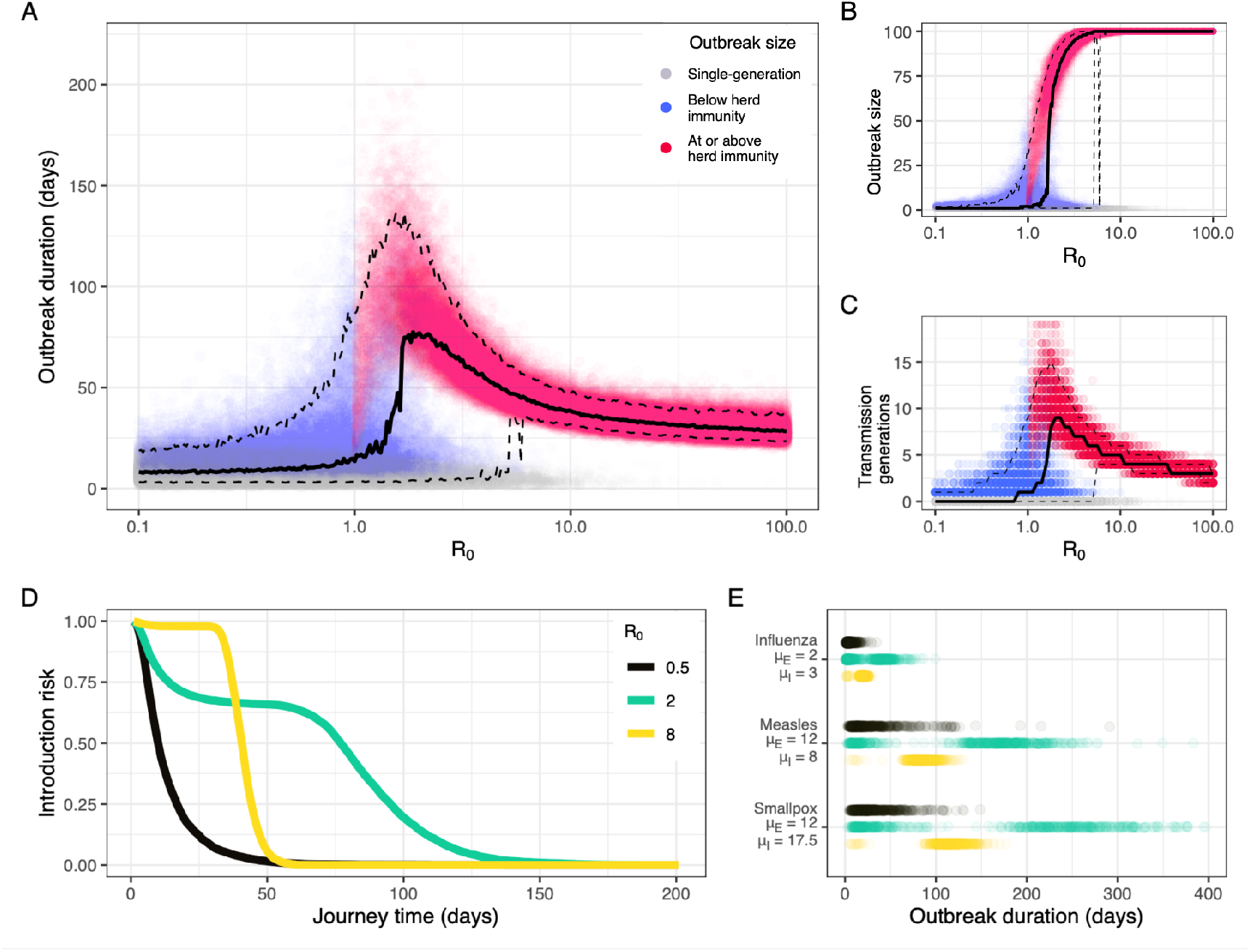
Basic Dynamics. (**A**) Outbreak duration, (**B**) outbreak size and (**C**) number of transmission generations by *R*_0_, assuming a fully-susceptible population of *N* = 100 and a theoretical pathogen with *µ*_*E*_ = *µ*_*I*_ = 5 days and *k*_*E*_ = *k*_*I*_ = 3. Solid black lines show median outbreak duration, outbreak size, and number of generations. Top and bottom dashed lines respectively show 95^th^ and 5^th^ percentile outbreak duration, outbreak size, and number of generations. (**D**) Probability of at least one person in state *E* or state *I* (“introduction risk”) for any given journey time by *R*_0_, using the same population and pathogen parameters. (**E**) Outbreak length distribution by *R*_0_ in a fully susceptible population of *N* = 100 for influenza, measles and smallpox, using epidemiological parameters detailed in Table S1.

Historical accounts indicate that transmission on board ships was substantially more intense than transmission in typical land settings (Text S1). Thus, we explore a broad range of transmission intensities. These are summarized by the epidemiological parameter *R*_0_, or the average number of infections that a single person will produce in a fully-susceptible population.

We observe three outbreak duration regimes, all of which depend heavily on transmission intensity. Under strongly subcritical transmission (*R*_0_ ≲ 0.8), the majority of simulations result in no transmission beyond the index case (Figure 1A). These “single-generation” outbreaks last only as long as the course of infection in a single person, in this case an average of 10 days.

Under strongly supercritical transmission (*R*_0_ ≳ 5), outbreaks are large and almost universally reach or exceed the threshold for ship herd immunity, 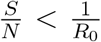 (Figure 1B). This reliably results in 35-55 day outbreaks for the modeled scenario, with duration steadily decreasing as *R*_0_ increases. Occasionally, simulations under strongly supercritical *R*_0_ also result in short single- or two-generation outbreaks (Figure 1A–1C); these are consistent with the occurrence of minor outbreaks due to random extinction in stochastic systems [39,40].

Values of *R*_0_ near criticality (0.8 ≲ *R*_0_ ≲ 5), produce the longest outbreaks, with median duration peaking around *R*_0_ = 2. These are made possible by extended, multigenerational transmission chains (Figure 1C). Yet while near-critical conditions give rise to the longest outbreaks, they also result in the widest range of outbreak durations. An *R*_0_ of 1 may result in outbreaks that last 150 days or more, but median outbreak duration under these conditions is just 14 days.

Transmission intensity modulates a pathogen’s overall introduction risk — here defined as the net probability that at least one passenger is carrying the pathogen (i.e. in state *E* or state *I*) upon arrival, for any given journey length. Under strongly subcritical transmission (e.g. *R*_0_ = 0.5), introduction risk decays rapidly with journey time, with 50% probability of introduction at 10 days and 25% probability at 17 days (Figure 1D). Under strongly supercritical transmission (*R*_0_ = 8), pathogen introduction risk is sigmoidal: introduction is near certain (*≥* 95%) for journeys of 33 days or less, then falls rapidly for journey times exceeding this threshold. Introduction is least predictable for weakly supercritical values of *R*_0_ (*R*_0_ = 2). Here, many outbreaks end quickly due to random extinction (Figure 1A). Yet past this threshold, risk broadly plateaus until much longer journey times (*∼* 60 days), then declines with a long tail. Thus, the relative introduction risk of weakly versus strongly supercritical transmission depends on journey length. Strongly supercritical transmission is significantly more likely to result in pathogen introduction for journeys of 33 days or less, since intense transmission carries minimal risk of early extinction. But across journeys of 40 days or more, pathogen introduction is most likely under weakly supercritical transmission.

For real pathogens, introduction thresholds are governed by pathogen-specific natural history, above all by the durations of a pathogen’s latent and infectious periods (Figure 1E). We explore outbreak length for influenza, measles, and smallpox at subcritical, near-critical, and strongly super-critical *R*_0_ (Table S1). The results demonstrate that relative introduction risk can be often inferred from pathogen natural history, even in the absence of shipboard *R*_0_ estimates. At any *R*_0_, smallpox typically survives longer on board a ship than measles, which in turn typically survives longer than influenza. Natural history also indicates some general introduction thresholds, which hold regardless of transmission intensity. For example, for a ship with 100 people on board, influenza introduction is extremely unlikely for journeys lasting longer than 100 days, regardless of *R*_0_.

### Incorporating Population Size and Susceptibility

Next, we expand our analysis beyond the unlikely scenario of one ship with *N* = 100 and 100% population-level susceptibility to consider the combined effects of ship population In populations with some initial immunity to infection (i.e. where 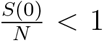 *<* 1), transmission intensity is most meaningfully measured as a pathogen’s “effective” reproduction number, *R*_*e*_. Because population immunity levels change over the course of an outbreak, this is commonly expressed as a function of time, i.e. *R*_*e*_(*t*). Notably, *R*_*e*_(*t*) is a linear function of a pathogen’s basic reproduction number, *R*_0_. Hence, 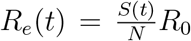, with critical transmission occurring at the threshold *R*_*e*_(*t*) = 1. We consider shipboard transmission at *t* = 0, where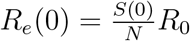.

First, we vary the total number of people who are initially susceptible, *S*(0), while holding *R*_*e*_(0) constant. We do so by fixing *N* = 1001, choosing *S*(0), and back-calculating *R*_0_ to maintain the same effective rate of transmission. This results in a roughly loglinear relationship between initial susceptible population size and outbreak duration at critical and supercritical values of *R*_*e*_(0) (Figure 2A). At *R*_*e*_(0) = 1, increasing *S*(0) has little influence on median outbreak duration but substantially increases 95th percentile outbreak duration. At *R*_*e*_(0) = 2, increasing *S*(0) increases both median and 95th percentile outbreak times. Finally, at *R*_*e*_(0) = 8, increasing *S*(0) dependably increases median, 5^th^ percentile, and 95^th^ percentile outbreak times.

**Fig. 2.**
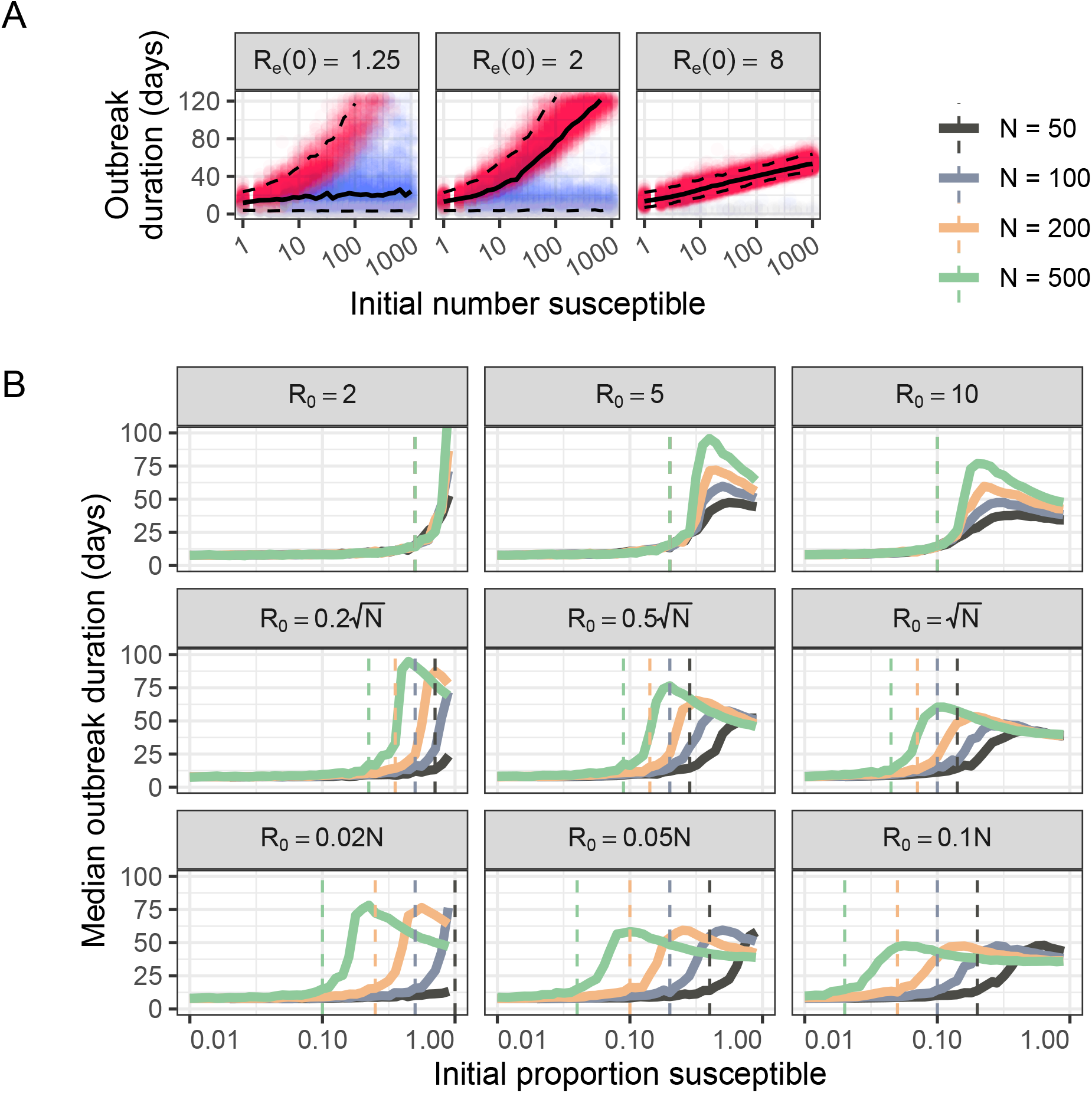
Effect of susceptible population size. (**A**) Outbreak duration by initial susceptible population (*S*(0)) and effective reproduction number (*R*_*e*_(0)). We fix *N* = 1001 and backcalculate *R*_0_ for each value of *S*(0) to maintain constant *R*_*e*_(0). As above, we base simulations on a theoretical pathogen with *µ*_*E*_ = *µ*_*I*_ = 5 days. Solid black lines show median outbreak duration; top and bottom dashed lines show 95^th^ and 5^th^ percentile durations respectively. Colors match those used in Figure 1A–1C and denote single-generation outbreaks (grey), outbreaks that reach herd immunity (red), and outbreaks that terminate before herd immunity is achieved (blue). (**B**) Median outbreak duration by ship size and by initial proportion susceptible. Rows show three density dependence scenarios: full frequency dependence (*q* = 0, top row); intermediate density dependence (*q* = 0.5, middle row); and full density dependence (*q* = 1, bottom row). Columns show three scenarios for transmission intensity: *β*_*fd*_ = 0.04 (left); *β*_*fd*_ = 1 (middle); and *β*_*fd*_ = 2 (right). Calculated using *µ*_*E*_ = *µ*_*I*_ = 5 days and *β*_*dd*_ = *β*_*fd*_*/*100 (Methods; Text S1) size, *N*, and initial proportion susceptible, 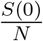, on ship outbreak duration.

Next, we vary *N* as well as *S*(0). This opens the question of what relationship we expect between *N, S* and *R*_0_ in the unique environment on board a historical ship. Records from the time indicate that many vessels suffered from inadequate ventilation and extreme levels of crowding (Text S1). On land, these conditions generally give rise to “density-dependent” patterns of transmission, where contact rates scale linearly with population size (*R*_0_ *∝ N*). Yet ships were also famously structured and compartmentalized environments, which typically align with assumptions of “frequency-dependent” transmission (Text S1); here contact rates are assumed to remain constant, regardless of total population size (*R*_0_ *⫫ N*).

In practice, we expect that effective density dependence varied substantially according to ship layout and construction, social norms, and pathogen-side biology. Thus we consider three density dependence scenarios: classical density dependence (*R*_0_ *∝ N*), classical frequency dependence (*R*_0_ *⫫ N*), and an intermediate degree of density dependence (*R*_0_ *∝ N* ^0.5^). Under each scenario, we explore the effect of initial ship susceptibility, 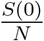, on median outbreak duration across several total population sizes, *N*.

In all circumstances, larger and more susceptible populations generally present greater risks of pathogen introduction across any given journey (Figure 2B). Yet they do so in different ways, and for different reasons.

Under classical frequency dependence, *R*_0_ = *µ*_*I*_*β*_*fd*_, where *µ*_*I*_ represents the average duration of an individual’s infectious period and where *β*_*fd*_ represents the average number of onward infections that a single infected person would generate, per day, in a fully susceptible population. Critical transmission, *R*_*e*_(0) = 1, occurs at the constant threshold 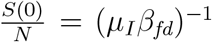. The threshold value of 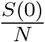 required for *R*_*e*_(0) = 1 is independent of total ship population, *N*. However, for any given 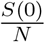, ships with greater *N* must have a proportionally greater number of susceptibles, *S*(0). Since ships with greater *S*(0) experience longer outbreaks at a given supercritical value of *R*_*e*_(0) (Figure 2A), ships with greater total populations display longer median outbreak times at *R*_*e*_(0) *>* 1 (Figure 2B, top row).

Under classical density dependence, *R*_0_ = *µ*_*I*_*β*_*dd*_*N*, where *β*_*dd*_ represents the average proportion of a given population, *N*, that a single infected person would infect per day in a fully-susceptible population. Critical transmission occurs at the threshold 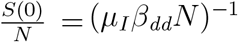. Multiplying both sides by a factor of *N* reveals that critical transmission depends solely on initial susceptible population size: *S*(0) = (*µ*_*I*_*β*_*dd*_)^*−*1^. When *N* is large, this threshold for *S*(0) represents a smaller fraction of the total population. But, in contrast to frequency-dependent transmission, *S*(0) is constant at any given *R*_*e*_(0), and so peak outbreak duration does not vary across ships of different sizes. Rather, larger ship populations give rise to near-critical and supercritical transmission at lower values of 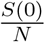 (Figure 2B, bottom row).

Under intermediate transmission, *R*_0_ = *µ*_*I*_(*β*_*fd*_*β*_*dd*_*N*)^0.5^. Critical transmission occurs at the threshold 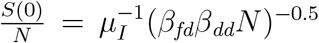, and at the total susceptibility level 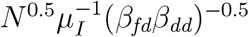. Under this model, larger ship populations reach critical transmission at slightly lower initial proportions of susceptibility, have a critical *S*(0) that scales sublinearly with *N*, and hence display slightly higher outbreak length for any given *R*_*e*_(0) (Figure 2B, middle row).

Finally, we note that regardless of density dependence, ships with a higher rate of contact (represented either by *β*_*fd*_ or by *β*_*dd*_) require lower initial susceptibility for critical transmission (compare across rows in Figure 2B). Thus, ships with higher rates of social mixing (e.g. more crowded ships) require fewer susceptible people to achieve supercritical transmission, regardless of total ship population size.

Thus, even in the absence of detailed reconstructions of ship transmission patterns, we can conclude that ships with larger, more crowded populations presented greater risks of pathogen introduction — be this by increasing total persistence times, decreasing the susceptibility fraction required for critical transmission, or both. In practice, the risk associated with larger ship populations was almost certainly boosted further by an increased chance of carrying at least one infected person on departure. We do not account for this difference, instead conditioning on the assumption that all ships depart with a single infected individual. Yet in cases with low infection prevalence at the port of origin, this elevated chance of having at least one infected person on board at the time of departure would have substantially increased net introduction risk. Thus, ships with larger populations were both more likely to depart with infection on board and more likely to sustain this infection outbreak until arrival.

### Historical Applications

Voyage characteristics such as journey time, population size, and population susceptibility varied substantially across different time periods, transit routes, and ship constructions. We explore some of this variation, and its implications, using port arrivals data for Gold Rush-era San Francisco, 1850-1852, originally collected by historian and genealogist Louis J. Rasmussen (Figure 3) [41–43].

**Fig. 3.**
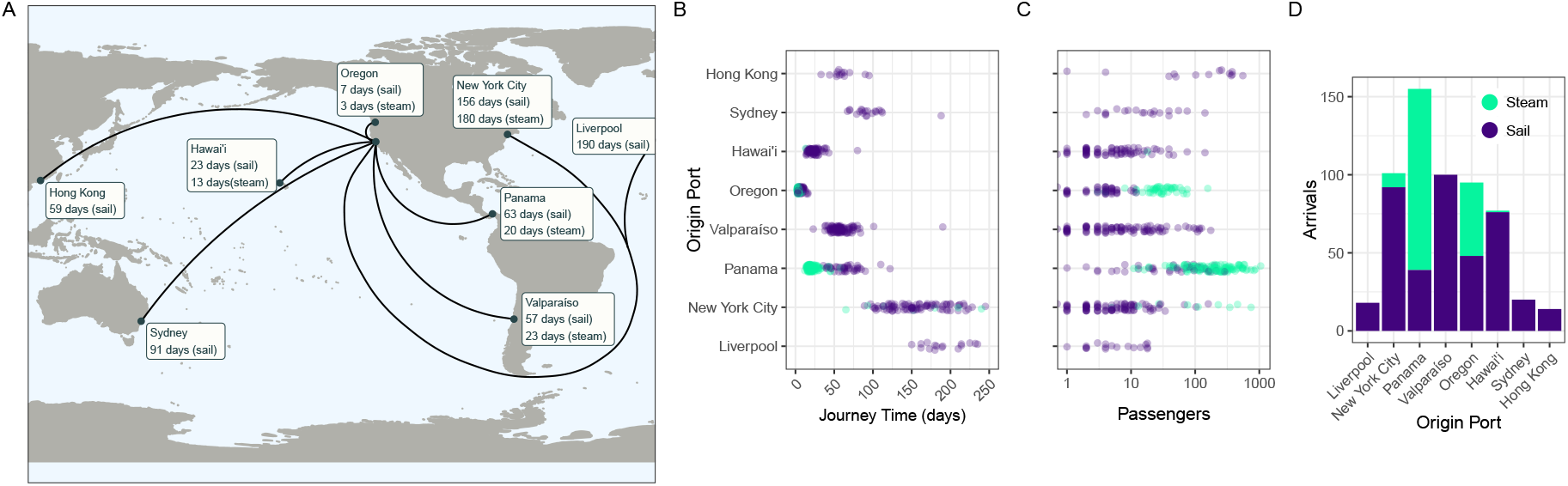
San Francisco Arrivals, June 1850 – June 1852. (**A**) Map of arrivals into San Francisco harbor, June 6^th^ 1850 June 9^th^ 1852, with median journey times by ship technology. (**B**) Journey time, (**C**) passenger number and (**D**) number of voyages by origin port and by ship technology. Data from Louis J. Rasmussen’s *San Francisco Passenger Lists* [41–43]

While acute respiratory infections first crossed the Atlantic ocean in the late fifteenth and early sixteenth centuries [13], by the mid-nineteenth century pathogens such as smallpox and measles had only recently begun to arrive across the Pacific basin. California saw its first region-wide outbreaks of smallpox and measles in 1806 and 1838, respectively [44–46]. Smallpox was first introduced to Australia in 1788 but did not see second introduction until 1829, while measles appears to have arrived for the first time in 1850 [36,47]. Several Pacific islands saw initial introductions well into the late nineteenth century, including Hawai’i (smallpox, 1853); Easter Island (smallpox, 1863); Fiji (measles, 1875); and Tonga (measles, 1893) [47].

During the years 1850-1852, passengers journeyed to San Francisco from across East Asia, Australasia, South America, and Europe. In an era preceding reliable transcontinental rail, ocean travel also provided one of the fastest and safest routes from eastern North America to the newly-established state of California [48]. Median sailing times ranged from 7 days (from Oregon Territory) to 190 days (from Liverpool, England), with considerable variation within routes (Figure 3A-B, Table S2). Longer-range sail voyages displayed an especially broad range of transit times. For example, sail journeys from New York City could be as long as 283 days (on the *Primoguet*) or as short as 89 days (on the *Flying Cloud* — reportedly “the fastest [sail] voyage ever recorded”) [42].

Steam travel represented a phase transition in transoceanic pathogen circulation for several reasons. First, in most cases, the technology dramatically reduced journey times. Median transit times from Panama were 63 days by sail but just 20 days by steam. Meanwhile, steam reduced median journey times from Oregon from 7 days to just 3 days (Table S2). These shorter journey times would have increased risk of shipborne pathogen introduction significantly.

Second, steam ships transported some of the greatest numbers of passengers (Figure 3C). Steamers from Panama carried a median of 196 passengers and as many as 1,050, compared with a median of 53 and a maximum of 287 by sail (Table S2). Oregon steamers carried a median of 28 passengers and as many as 157, compared with a median of 4 and a maximum of just 12 by sail. Finally, steam vessels from New York City carried a median of 111 passengers and a maximum of 743, compared to a median of 5 and a maximum of 160 by sail. The only sail route that could compete with steam travel on passenger numbers was the route from Hong Kong, which transported a median of 201 passengers and a maximum of 553. As demonstrated above, larger passenger numbers would have substantially increased ships’ capacity for sustained pathogen circulation.

Finally, steam travel represented some of the most frequent voyages (Figure 3C), resulting in a greater cumulative risk of pathogen introduction across any given period. Particularly striking are the 116 steam journeys from Panama that arrived between June 1850 and June 1852.

To explore possible differences in introduction risk across each route and type of ship, we simulate influenza, measles, and smallpox outbreaks across the full range of ship populations and journey times represented in the San Francisco dataset (Figure 4A). Here, contours represent pathogen introduction risk by journey time and by total ship population, assuming 5% population-level susceptibility and intermediate density dependence, and calibrating transmission intensity with reference to standard literature values and analyses of shipboard outbreaks (Text S1, Table S1). We overplot individual journeys into San Francisco to assess pathogen introduction risk across each route (Figure 4A). We plot a selection of routes on each panel for visual clarity. However, the observations below concern introduction risk for all pathogens across all routes travelled.

**Fig. 4.**
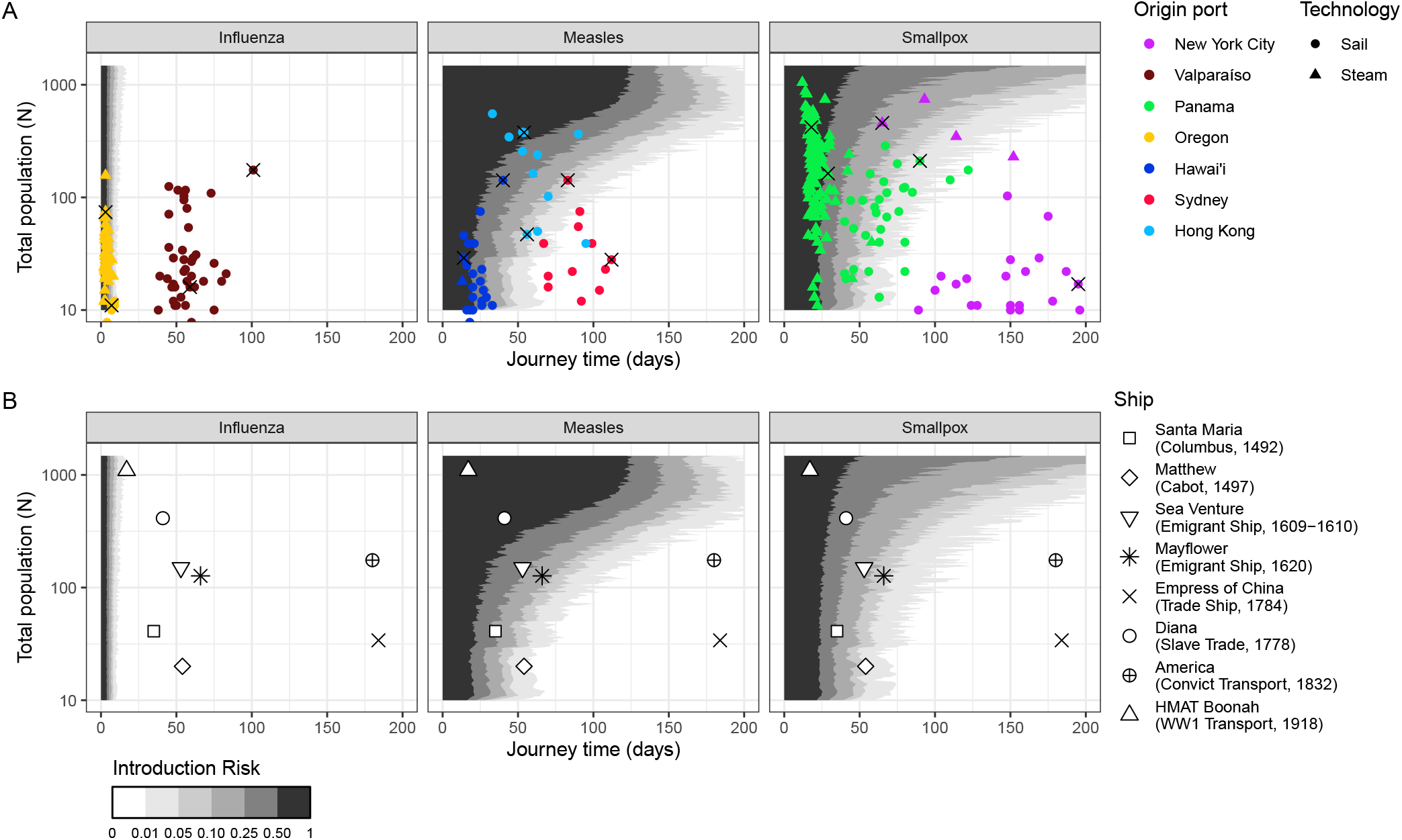
Historical Applications. Introduction risk for influenza, measles, and smallpox by journey time and by total ship population, *N*, assuming 5% initial population-level susceptibility, intermediate density dependence (*q* = 0.5), and *µ*_*E*_, *µ*_*I*_, and *β*_*fd*_ according to consensus natural history parameters (Table S1). We back-calculate *β*_*fd*_ as 1*/µ*_*I*_ times a pathogen’s typical land *R*_0_, and set *β*_*dd*_ = *β*_*fd*_*/*75 (Text S1).(**A**) overplots data on San Francisco Port arrivals, June 1850 – June 1852. Here, total population (*N*) represents only the passengers on board each ship, as crew data is not available. Introduction risks for all three pathogens are shown for 16 selected voyages in Table 1; these voyages are indicated with black crosses. For the two ships with documented infectious disease outbreaks, the *Gold Hunter* and the *Sir Charles Napier*, we also performed sensitivity analyses investigating the robustness of our predictions to different rates of transmission and population-level susceptibility (Figure S1). (**B**) overplots selected historical journeys, 1492–1918, chosen to be indicative of the broad trends in transoceanic shipping. *N* represents the combined totals of passengers and crew. Sources and further data are available in Table S3. Numerical introduction risk estimates for (**B**) are provided in Table 2.

For select voyages we also provide numerical introduction risk estimates for each pathogen in Table 1. Particularly interesting are two voyages from Panama with documented outbreaks of acute viral infections: the *Gold Hunter* steam ship, which arrived in San Francisco after a 29-day voyage with one active case of smallpox, and the *Sir Charles Napier* sail ship, which experienced an outbreak of “measles, dysentery, and fever” lasting “about three weeks” of its ninety-day voyage.

**Table 1.** Numerical introduction risk estimates for influenza, measles, and smallpox across selected voyages to San Francisco, 1850-1852.

| From | Type | Days | N | Pathogen Introduction Risk |  |  |
| --- | --- | --- | --- | --- | --- | --- |
|  |  |  |  | Influenza | Measles | Smallpox |
| New York | Sail | 195 | 17 | < 0.001 | < 0.001 | < 0.001 |
| New York | Steam | 65 | 458 | < 0.001 | 0.611 | 0.207 |
| Valparaíso | Sail | 59 | 16 | < 0.001 | 0.010 | 0.036 |
| Valparaíso | Sail | 101 | 175 | < 0.001 | 0.053 | 0.013 |
| Panama* | Sail | 90 | 211 | < 0.001 | 0.160 | 0.057 |
| Panama† | Steam | 29 | 163 | < 0.001 | 0.556 | 0.422 |
| Panama | Steam | 18 | 420 | 0.001 | 0.813 | 0.749 |
| Oregon | Steam | 3 | 74 | 0.656 | 0.996 | > 0.999 |
| Oregon | Sail | 7 | 11 | 0.095 | 0.931 | 0.979 |
| Hawai'i | Sail | 40 | 142 | < 0.001 | 0.399 | 0.254 |
| Hawai'i | Sail | 14 | 29 | 0.001 | 0.692 | 0.817 |
| Sydney | Sail | 112 | 28 | < 0.001 | < 0.001 | < 0.001 |
| Sydney | Sail | 83 | 142 | < 0.001 | 0.076 | 0.032 |
| Hong Kong | Sail | 56 | 47 | < 0.001 | 0.040 | 0.047 |
| Hong Kong | Sail | 54 | 377 | < 0.001 | 0.613 | 0.224 |

\* This vessel, the *Sir Charles Napier*, experienced an outbreak of “measles, dysentery, and fever” which lasted “about three weeks”. 36 passengers died, with the final death occurring 54 days into the voyage [49] (Table S4; Figure S1).
† This vessel, the *Gold Hunter*, arrived in San Francisco with one active smallpox case. The patient was isolated on arrival [50]. (Table S4; Figure S1)

**Table 2.** Numerical introduction risk estimates for influenza, measles, and smallpox across selected historical voyages, 1492-1918.

| Vessel | Days | N | Pathogen Introduction Risk |  |  |
| --- | --- | --- | --- | --- | --- |
|  |  |  | Influenza | Measles | Smallpox |
| *Santa María* | 35 | 41 | < 0.001 | 0.242 | 0.235 |
| *Matthew* | 54 | 20 | < 0.001 | 0.021 | 0.044 |
| *Sea Venture* | 53 | 150 | < 0.001 | 0.318 | 0.160 |
| *Mayflower* | 66 | 127 | < 0.001 | 0.145 | 0.073 |
| *Diana* | 41 | 443 | < 0.001 | 0.667 | 0.345 |
| *Empress of China* | 184 | 34 | < 0.001 | < 0.001 | < 0.001 |
| *America* | 180 | 175 | < 0.001 | < 0.001 | < 0.001 |
| HMAT *Boonah* | 17 | 1095 | 0.005 | 0.905 | 0.823 |

Influenza’s relatively low *R*_0_ and extremely fast generation period result in a very low risk of introduction into San Francisco from any origin port except Oregon and perhaps Panama and Hawai’i. Even then, only the fastest voyages presented any significant risk of pathogen transfer. Had a person with influenza been present on board the *Columbia* steam ship (3 days, 74 passengers) at its time of departure from Oregon, we estimate a 66% risk of introduction into San Francisco (Table 1). By contrast, we estimate just a 1% risk for the *Tarquina* sail ship from Oregon (7 days, 11 passengers), a 0.1% risk on the *Columbus* steam ship from Panama (18 days, 420 passengers) and a 0.1% risk on the *Baltimore* sail ship from Hawai’i (14 days, 29 passengers).

Measles, with longer latent and infectious periods, presents moderate introduction risks across all journey times ≲ 40 days (Table 1) – consistent with this pathogen’s range of durations for single-generation outbreaks (Figure 1E) This range includes the vast majority of journeys originating from Oregon (by steam or sail), Hawai’i (by steam or sail), and Panama (by steam). Additionally, we estimate plausible introductions from Panama (by sail) and Hong Kong, especially on ships transporting large populations. Had the *Iowa* sail ship (54 days, 377 people) departed Hong Kong with a measles patient on board, we estimate a 61% chance of introduction despite the long journey (Table 1). Had the *Golden Gate* steam ship (65 days, 458 passengers) departed New York City with one infected passenger, we likewise estimate a 61% introduction risk. Our results are also consistent with reports that the *Sir Charles Napier* sail ship from Panama experienced an outbreak of measles which ended roughly 36 days before arrival; under our base assumptions, we estimate that this vessel had just a 16% chance of sustaining the pathogen across the duration of its 90-day voyage. Supplementary analyses indicate measles introduction following this voyage would have been plausible under some circumstances, particularly if a higher proportion of the population had been susceptible and if shipboard transmission intensity had intermediate intensity (Figure S1).

Smallpox has a substantially longer generation period than either measles or influenza (*µ*_*E*_ = 12 days; *µ*_*I*_ = 17.5 days). Consequently, journeys of ≲ 50 days present a moderate introduction risk at any ship population size, mirroring the introduction range described above for measles. As before, we estimate higher introduction risks for ships with larger populations (Table 1). Yet since smallpox is less transmissible than measles, ships require larger population sizes to achieve a equivalent *R*_*e*_(0). Thus, in some cases highly-populated ships present a lower risk of introducing smallpox than they did measles. For instance, had the *Golden Gate* departed New York City with one infected patient on board, we estimate just a 21% risk of smallpox introduction (Table 1). Our results are consistent with the one documented introduction of smallpox by ship to San Francisco during our period of study: under base assumptions, we estimate that the *Gold Hunter* had a 42% chance of arriving with at least one active case. Supplementary analyses indicate that smallpox introduction following this vessel’s 29-day voyage was in fact highly likely across a wide range of conditions, even including scenarios with no susceptible people on board besides the index case (Figure S1). This is intuitive; given the pathogen’s lengthy latent and infectious periods (we assume mean values of 12 and 17.5 days, respectively), a single case could easily last as long as the *Gold Hunter* ‘s period in transit (Table S1).

Finally, we use these analyses to inform the plausibility of ship-borne pathogen transfer across a selection of historical voyages, chosen to reflect the variety of shipping routes, technologies, and practices between the 15^th^ and 20^th^ centuries (Figure 4, Table S3). For these analyses we again assume a 5% rate of susceptibility, although in practice we expect this rate varied significantly by location, time period and ship population.

Under these assumptions, early transatlantic voyages of exploration could plausibly have introduced measles or smallpox to their places of arrival (Figure 4B, Table 2). We estimate a 24% chance of measles introduction and an equal chance of smallpox introduction had Christopher Columbus’s 1492 voyage on *Santa María* (35 days, 41 people) departed with one case of either pathogen on board. We estimate a 2% risk of measles introduction and 4% risk of smallpox introduction on John Cabot’s 1497 exploration on the *Matthew* (54 days, 20 people). Introduction risks for both pathogens were substantially higher on the transatlantic slave trade ship *Diana*, which carried 443 enslaved people and crew from Îles del Los, off the coast of West Africa, to Curaçao, in the Caribbean: a 67% risk for measles and a 35% risk for smallpox, had one person been infected at the time of departure.

Meanwhile, the lengthy journey times of the *Empress of China* trade ship (184 days), which traveled from New York City to present-day Macao, and the *America* convict ship (180 days), which traveled from the United Kingdom to Australia, suggest a compelling explanation for the substantially later introduction of smallpox and measles to the South Pacific. Even with 175 passengers, a voyage such as the *America*’s is outside the range of plausible introduction for all three pathogens.

Our analyses indicate that by far the greatest introduction risk of smallpox and measles – and the only plausible influenza introduction – came from fast, highly-populated ships such as the WW1 troop ship HMAT *Boonah* (1,095 passengers and crew), here undertaking a 17-day journey from South Africa to Australia. Had this ship departed with one infected person on board, it would have had a 0.5% risk of introducing influenza, an 82% chance of introducting measles and a 91% chance of introducing smallpox to its destination. This combination of fast transit and extremely large passenger populations substantially increased both the magnitude of introduction risk for moderately fast-burning pathogens (such as measles and smallpox) and expanded the range of potential introduction to include pathogens (such as influenza) with much faster life cycles.

## Discussion

Many stories of transoceanic pathogen transfer have focused heavily on early colonial European seafaring. Our analysis indicates that introductions of smallpox and, to a lesser extent, measles from Europe to the Americas via early colonial voyages was plausible, but by no means guaranteed. Depending on weather, these journeys could last just 5–10 weeks [51], which is a reasonable time frame for these pathogens to persist on board a ship. In these contexts, overall pathogen introduction rates likely depended more on population-side factors – for example, the density of susceptible people, or the rate at which ships departed with active infection(s) on board – than on the precise epidemiological parameters aboard ships. By contrast, our model shows that early transatlantic introductions of faster-burning pathogens such as influenza were unlikely, as were introductions of any acute pathogen on longer journeys such as sail voyages to the Pacific [32].

More recently, the story of transoceanic pathogen transfer has been told as one of technological innovation. Our work supports and extends Cliff and Haggett [33]’s argument that steam technology transformed rates of transoceanic pathogen transfer. Steam ships travelled more quickly, could carry greater numbers of passengers and, in the case of Gold Rush-era San Francisco, made more frequent voyages. Under the right conditions, this could have increased both the rate and the geographic range of transoceanic pathogen transfer substantially (Figure 4A; Table 1).

However, steam travel was not unique in enabling global pathogen circulation. Our analysis confirms longstanding arguments by historians that processes which involved large-scale people-movement—for example war, migration, or the transatlantic slave trade—were enormously significant for global pathogen ecology [8,9,52,53]. In the case of 1850s California, ship population size could easily have been the difference between plausible introduction and epidemiological isolation. In 1852, two ships sailed from Hong Kong, the *Catalpa* and the *Iowa*. Both displayed similar transit times into San Francisco: 60 days and 54 days, respectively [41,42]. Yet while the *Catalpa* carried “Chinese merchandise, rice, cordage, and assorted goods” – along with one solitary passenger – the *Iowa* brought “377 unidentified in steerage”, likely Chinese people bound for California’s gold fields [54]. As our analyses show, the presence of 377 people on board transformed the *Iowa*’s capacity to sustain outbreaks of smallpox and measles across the journey from Hong Kong to San Francisco (Figure 4A; Table 1).

Our study specifically considers a small subset of human pathogens, chosen both for their historical impact and because they permit simple modeling approaches. Historical scholarship, together with recent advances in paleogenomic sequencing, demonstrates that transoceanic shipping enabled the diffusion of a much broader range of diseases [12,55,56]. These include pathogens with food-, water- and fomite-borne transmission (e.g. cholera, *Salmonella*) [55,57]; pathogens with vector-borne transmission (e.g. malaria, yellow fever, West Nile virus) [12,56,58–60]; pathogens with multi-species transmission (e.g. plague, tuberculosis) [56,61,62]; and pathogens which infected only non-human animals (e.g. rinderpest, foot-and-mouth disease) [63,64]. Transoceanic shipping also shaped the global dissemination of broad range of plant and animal species; recent scholarship suggests that these processes were likewise shaped by the speed and volume of transoceanic shipping, as well by trade of specific commodities [65–68]. A full understanding of these introductions will require modified modeling approaches and likely additional historical data. This issue is also pertinent to smallpox, for which the extent of fomite transmission is unclear. Recent research indicates that orthopoxviruses can remain viable on surfaces for weeks [69]. The World Health Organization’s smallpox eradication campaign found that fomites caused only a small minority of outbreaks [70], but in the context of historical pathogen circulation even rare introductions can be impactful [28–30].

Several additional questions require further consideration. One concerns the mechanics of shipboard transmission. Little is known concerning either the density dependence or the intensity of transmission on board historical vessels (Text S1). Our analysis points to several strong qualitative patterns, which are robust across a broad range of parameters. Crowded ships with larger and more susceptible populations presented greater risks regardless of the precise form of density dependence (Figure 2). Similarly, broad ranges for plausible outbreak duration can be inferred from pathogen natural history, even without knowing shipboard *R*_0_ (Figure 1E). More refined quantitative predictions will depend on the specifics of particular ships and voyages, and will require further research into shipboard transmission dynamics. We have used our model to map the possible effects of assumptions regarding density dependence and transmission intensity across a range of plausible parameters (Figure S1–Figure S4).

A second question concerns the extent of shipboard population structure. Ships were famously hierarchical environments, and highly compartmentalized populations may have prolonged ship outbreaks. Captains or surgeons may also have manipulated population structure in response to outbreaks, for example through case isolation, quarantine, or disembarkation of known or suspected infections. While population structure almost certainly shaped outbreak duration, incorporating these effects is challenging in the absence of high-resolution outbreak data from a given ship. Moreover, on ships with poor ventilation or hygiene practices, transmission could plausibly have been homogeneous regardless of social behaviours or most medical interventions.

Our work sheds light on how shipboard transmission dynamics shape introduction risk, but reconstructing historically accurate circulation rates would require more information regarding pathogen dynamics in source populations. This matters for inferring likely immunity rates in ship populations, which our model shows can have a large impact on estimated risks (Figure S5). It also matters for assessing the probability of at least one infected individual on board ship at the point of departure. Longitudinal mortality data exists for diseases such as smallpox and measles, especially in European and North American contexts, with particularly well-preserved time series in the London Bills of Mortality [27,71]. Reconstructing historical prevalence and immunity landscapes from these sources is difficult, but is critical for estimating realistic pathogen transfer rates pre-20^th^ century contexts.

A related question concerns the contribution of partly-immune individuals to pathogen circulation within a given population. The ability of partly-immune people to be infected and transmit infection has long been recognized as an important driver of influenza [72] and smallpox [70] epidemiology. Partial immunity also provides a compelling explanation for recent resurgences in mumps [73] and pertussis [74,75]. Moreover, it is plausible that the contribution to transmission from partly-immune individuals was more significant on board a ship than it was on land, due to extended exposures or large infectious doses. This possibility – and the influence of partial immunity on outbreak duration more broadly – require further investigation.

Our model offers a general assessment of outbreak duration in a closed population, which holds significance beyond historical systems. Understanding infection persistence in discrete subpopulations is critical for studying pathogen circulation in any system with limited host connectivity, from wildlife populations [76,77] to agricultural biosecurity [78,79] to human populations distributed across regions [17,80].

These findings carry important historical implications, connect to present-day disease dynamics, and may, some day, inform interplanetary risk of pathogen spread. Centuries before the present-day upheaval of air travel [80–82] and large-scale human re-locations [83] combined to transform global pathogen ecology. Yet this process was almost certainly lengthy, geographically uneven, and contingent on complex interplays between technology, shipping practices, and specific pathogen biology. This presents a rich avenue for collaboration between ecologists, epidemiologists, historians, and social scientists. How do social, economic and technological forces combine to shape global pathogen ecology – and with what consequences along the way for the world’s people, places, and pathogens?

## Materials and Methods

### Model Description

We simulate shipboard outbreaks using a stochastic SEIR model (Text S3). We implement continuous-time stochastic simulations in R with the Gillespie Algorithm, using the package GillespieSSA [84]. All simulations assume a single index case in state *E* at the time of departure. We define outbreak duration as the time until both state *E* and state *I* contain zero individuals.

To achieve a more realistic depiction of the time course of infection, we use the Linear Chain Trick to make dwell times in state *E* and state *I* Erlang-distributed [85]. For all simulations, we use shapes *k*_*E*_ = *k*_*I*_ = 3 and rates *k*_*E*_*/µ*_*E*_ and *k*_*I*_*/µ*_*I*_ for states *E* and *I* respectively. This technique gives a unimodal distribution with a long right-hand tail, such that disease progression is relatively constrained in most individuals, but occasional individuals experience substantially longer periods of incubation or infectiousness [86]. We assume that state *E* is pre-symptomatic and that initially infected individuals could board ship at any point during this period, randomly assigning index cases across substates *E*_1_, *E*_2_, …, *E*_*kE*_ at the point of departure.

Our model also tracks infection across pathogen generations, when needed. The *I*_*n*_ infectious individuals from generation *n* produce new exposed individuals *E*_*n*+1_, which represent the (*n* + 1)^st^ generation of infections.

To account for uncertainty and variation in the density dependence of shipboard contact rates, our model uses a flexible depiction of density dependence encoded by the equation:

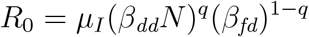

*R*_0_ is the pathogen’s basic reproduction number on board a given ship. This represents the average number of infections that an infected person generates in a fully-susceptible population, where *µ*_*I*_ is the average period of infectiousness. The density dependence of transmission is adjusted with the parameter *q*, with *q* = 1 representing classical density-dependent transmission (*R*_0_ *∝ N*), *q* = 0 representing classical frequency-dependent transmission (*R*_0_ *⫫ N*), and 0 *< q <* 1 representing intermediate density dependence (*R*_0_ *∝ N* ^*q*^). The parameters *β*_*dd*_ and *β*_*fd*_ modulate the intensity of transmission under each density dependence pole — intuitively, the proportion (*β*_*dd*_) and the raw number (*β*_*fd*_) of people on board ship that a single infected individual will infect per day, on average, in a fully-susceptible population.

In analyses where *N* is constant and where we do not explore the effect of density dependence, we set *q* = 1 such that *R*_0_ = *β*_*dd*_*N*; we then back-calculate *β*_*dd*_ from *R*_0_ and *N*. Mathematically, this is equivalent to setting *q* = 0 with *β*_*fd*_ fixed at *β*_*fd*_ = *β*_*dd*_*N*.

For analyses where N varies, we infer *β*_*fd*_ from literature values of *R*_0_ and *µ*_*I*_ (Table S1) and set *β*_*dd*_ = *β*_*fd*_*/c*. Here, *c* is a constant representing the population size at which *β*_*dd*_ and *β*_*fd*_ would be equal. In main text analyses, we set *c* = 100. We explore alternative values of *c*, as well as a range of *q* values, in Figs. S2–S4.

### Historical Data

To provide real-life context for our theoretical results, we collected data on ship arrivals into the port of San Francisco between June 6 1850 and June 19 1852 from volumes I, II and III of genealogist and historian Louis. J. Rasmussen’s reference book *San Francisco Ship Passenger Lists* [41–43]. We recorded the port of origin, the ship type, the journey time, and the number of passengers for ships originating from seven locations: Hawai’i; Hong Kong; Oregon Territory; New York City; Sydney, Australia; Valparaíso, Chile; and Liverpool, England. In the few cases where Rasmussen reports ships making multiple stops at one or more of these locations in the course of their voyage, we record both the journey time into San Francisco from a ship’s port of origin and, where available, journey times into San Francisco from the intermediate port(s). We exclude ships where substantial numbers of people (*N >* 10) boarded during a voyage, as our model does not account for changes in population size subsequent to the initial point of departure.

For almost all ships, Rasmussen provides passenger numbers but not numbers of crew. We assume that in most cases, crew (i) represented a small proportion of a ship’s total population; (ii) were, as professional sailors, more likely to possess immunity to common maritime infections, and so represented an even smaller proportion of a ship’s susceptible people. Thus, in the absence of crew size data, analyses considering population size of vessels arriving into San Francisco approximate *N* as the total number of passengers on board each ship.

## Author Contributions

EB: conceptualization, data curation, formal analysis, investigation, methodology, visualization, writing – original draft, writing – reviewing/editing. JLS: conceptualization, formal analysis, funding acquisition, methodology, supervision, writing – reviewing/editing.

## Acknowledgments

We thank Benjamin Madley, Daniel Blumstein, Brandon Ogbunu, the Lloyd-Smith Lab, and two anonymous reviewers for their feedback on this work. Portions of the paper were developed from the thesis of EB.

## Supplementary Material

### Text S1. Transmission on board Historical Ships

#### Historical Evidence

##### Intensity

We expect that transmission on board historical ships was substantially more intense than transmission in land settings. In a study of pandemic 1918 influenza, Vynnycky et al. [87] estimated *R*_0_ values of approximately 4-11 on board the troop ship His Majesty’s Australian Transit (HMAT) *Boonah*, approximately 5-17 on board the troop ship HMAT *Devon*, and approximately 3.5-8 on board the troop ship HMAT *Medic*. These compare to an estimated range of roughly 1.5-4 in American and Scandinavian towns and cities. Similarly, White and Pagano [88] calculated an *R*_0_ of roughly 4.98 for the same epidemics on board HMAT *Boonah* and HMAT *Medic*, compared with an *R*_0_ of 1.34-3.21 in Maryland communities.

Similar analyses are not available for earlier time periods. However qualitative descriptions likewise indicate that conditions on board pre-1918 ships were highly conducive to intense pathogen transmission. One 1801 newspaper report describes an emigrant ship from Ireland to New York City so crowded that ‘the space between decks, occupied by nearly 300 persons, became the receptacle of all excremental matters’. [89]. Half a century later, English author Frank Marryat recalled travelling from Panama to San Francisco on a boat “so crowded with passengers, that it was not until it was ascertained that there was scarcely standing-room for those on board that she tripped her anchor” [90]. Longer excerpts from both texts, printed below, offer vivid depictions of the conditions that passengers suffered during these voyages.

In addition to crowding, pre-1918 ships also faced substantial ventilation challenges. On sail ships, opening portholes risked losing heat and letting in water. Thus, below-deck spaces were sealed, resulting in notoriously poor air quality [91,92]. The steam revolution somewhat improved onboard environments, both by providing a source of heat and, in some cases, powering active ventilation systems. Yet steam ventilation still failed – sometimes catastrophically. In 1848, seventy-three people died from suffocation on board the *Londonderry* emigrant steamer when hatches were closed during a storm [92]. Six years later, fifty passengers out of a total of 204 died by suffocation in an unnamed ship carrying emigrants from Mauritius [93]. “Under no circumstance can a ship of any kind be made as healthful or as comfortable as the house on shore,” wrote United States Navy surgeon Albert L. Gihon in 1886. Gihon added: “It is, practically, a floating box sealed against the admission of water, and, of course, also of air.” [94]

##### Density Dependence

The nature of density dependence of pathogen transmission on board historical ships is unclear. On land, early-twentieth century studies demonstrate a clear and intuitive link between population density and rates of respiratory infection. Notably, Brewer [95] documented higher rates of pandemic influenza in military units training in more crowded environments in a 1918 study of Fort Humphreys, Virginia.

Some contemporaries clearly thought of ship transmission in what we would today consider density-dependent terms. In a report on 1918 pandemic influenza on board Royal Navy ships, surgeon-commander Sheldon F. Dudley argued that ‘infective material must become so dense and diffused as to saturate the ship. That is to say, everyone on board receives a dose of the specific agent sufficient to cause influenza’ [96]. Dudley’s report is excerpted below.

Yet density dependence does not appear to describe every instance of shipboard transmission. In an analysis of smallpox outbreaks on board vessels bound into Australia, 1850-1908, quarantine director J. H. Cumpston found that most shipboard smallpox transmission was limited to close contacts of infected patients, such as family members, close colleagues, or those sharing beds and cabins [97]. Such a pattern typically argues for ‘frequency-dependent’ transmission, in which an infected individual on average transmits only to a fixed number of close contacts, regardless of a ship’s total population size.

##### Contemporary Examples

Even with the benefits of modern hygiene, sophisticated air circulation, and more highly-regulated living conditions, infectious disease outbreaks are common on present-day cruise [98–100], cargo [101] and naval ships [102]. Attack rates are often high. Vera et al. [103] reported a 49.1% attack rate of 2009 H1N1 pandemic influenza in an unnamed Peruvian Navy ship carrying 355 people, with greatest risk in cadets assigned high-density living quarters. Earhart et al. [104] observed a 42% attack rate of H3N2 influenza across more than 500 people on board Navy vessel USS *Arkansas*, despite 95% vaccination coverage. Brotherton et al. (2000) reported a 37% attack rate of influenza-like illness on a cruise ship carrying over 1600 passengers and crew.

Recent reports of SARS-CoV-2 transmission on board passenger ships [105–108] and military ships [109] indicate high attack rates across both vaccinated and unvaccinated populations. Multiple studies suggest that outbreaks could spread broadly across ship populations. Using a mechanistic model, Azimi et al. [105] estimate similar transmission contributions from long-range aerosols and from short-range aerosols and droplets on board the *Diamond Princess* cruise ship. Meanwhile, in a study of an extensive outbreak on board a Dutch river cruise (60 of 132 passengers infected), Veenstra et al. [108] observed no apparent clustering by cabin layout or by meal-time seating arrangements.

#### Density dependence parameter choices for historical analyses

Given the limited quantitative studies of density dependence on board historical ships, parameter choices for the analyses supporting Figure 4, Table 1, and Table 2 are necessarily approximations. We assume intermediate density-dependence (*q* = 0.5) and infer *β*_*fd*_ from standard literature values for *R*_0_ and for each pathogen’s infectious period, *µ*_*I*_ (*β*_*fd*_ = *R*_0_*/µ*_*I*_) (Table S1). We set *β*_*dd*_ = *β*_*fd*_*/c*, where *c* intuitively represents the value of *N* at which *β*_*fd*_ and *β*_*dd*_ would be equal. Using the general expression *R*_0_ = *µ*_*I*_(*β*_*dd*_*N*)^*q*^(*β*_*fd*_)^1*−q*^, and substituting *β*_*dd*_ = *β*_*fd*_*/c*, we obtain *R*_0_ = *µ*_*I*_*β*_*fd*_(*N/c*)^*q*^. Thus for our assumption of *q* = 0.5 on a ship, the ratio of shipboard *R*_0_ to frequency-dependent transmission (*R*_0_ = *µ*_*I*_*β*_*fd*_) is (*N/c*)^0.5^. To choose an appropriate value of *c*, we referred to the analyses of shipborne *R*_0_ by Vynnycky et al. [87] and White and Pagano [88], discussed above. In ship settings, Vynnycky et al. [87] estimate an *R*_0_ of *∼*4–17 for 1918 influenza on three *∼*1,000-person ships, compared with an *R*_0_ of 3.5–8 on land. White and Pagano [88] re-analysed two of the three ships in Vynnycky et al.’s analysis and found an *R*_0_ of *∼*4.97, compared with an *R*_0_ of 1.34–3.21 on land. Altogether, these analyses suggest that *R*_0_ for influenza on board a 1,000-person ship is roughly 2.5to 4.5-fold greater than its value on land. Assuming that these land *R*_0_ values represent frequency-dependent transmission, and assuming intermediate density dependence (*q* = 0.5) on board the three ships in these analyses, we back-calculate a *c* range of roughly 50–150. We use *c* = 100 in main text analyses. We explore additional values (*c* = 50, *c* = 150) in supplementary figures S2–S4, as well as a broader range of *q* values.

### Text S2. Qualitative Descriptions of Shipboard Infection and Transmission, 1801-1921

Three written descriptions of disease transmission on board nineteenth- and early twentieth-century ships are excerpted below. We do not intend these extracts to give a comprehensive view of shipboard transmission, but rather to offer a qualitative view of possible transmission scenarios. All spelling and grammar is original.

#### “Another instance of pestilence engendered in a ship crowded with passengers from Ireland.” [89]

The ship Nancy, Capt. John Herron, was charterd by a commercial house at Sligo, to carry passengers from that port to New-York. She sailed from Sligo on the 12th July, 1801, and arrived, after a passage of 77 days, at the port of New York, on the 27th of September following. This ship, of the burthen of 202 tons, received on board 417 passengers, and was navigated by nine seamen. The provisions, mere refuse, put up by government-contractors with the view of saving expense, were of the worst kind: and the water, which was also of bad quality, from the unexpected length of the voyage, became extremely scanty before the arrival of the ship.

In order to receive so great a number of passengers on board of this ship, temporary cabbins [sic] were built on the quarter-deck, which were filled with eighty persons. Three hundred were crowded into the space between decks.

It will excite no surprise that a vessel thus crowded became sickly soon after sailing from Sligo. Typhous fever and dysentery began to prevail, and destroyed the lives of a large proportion of the passengers.

In addition to the wretchedness of being confined in so small a space, these unhappy emigrants suffered all the evils which their habits of uncleanliness could produce. Their bodies and clothes, covered and saturated with filth, exhuded poison all around them. Partly from the want of strength and assistance among the sick, and partly from the want of a sense of decency, the space between decks, occupied by nearly 300 persons, became the receptacle of all excremental matters, insomuch that they issued in streams from the scuppers. The filth on the upper deck was nearly over the shoes. The sides of the ship were daubed and incrusted [sic] with excrements; and even the rope for the support of such that wished to go on board were unfit to be handled. The stench was intolerably offensive.

In such condition arrived this unfortunate vessel at the place assigned for quarantine in the port of New York. Ninety persons had died on the passage; one hundred and eighty were sick. Scarcely a healthy countenance was to be seen on board of the ship; very few had escaped disease; and many had suffered from three to four relapses. About forty were taken ill after their arrival.

As soon as possible after their arrival the sick were brought ashore; stropped of their filthy and pestilential clothes; their bodies thoroughly washed and scoured with soap and water; and then wrapped up in clean blankets, and carried into the wards appointed for their reception in the Marine Hospital. The permanent buildings of the establishment were insufficient to receive so great a number; tents, and other temporary accommodations, were provided for the remainder. Separation, ventilation, and cleanliness, as soon as they could be brought into action, accomplished every thing that could be expected. And only twenty-six have died since their arrival at this port.

#### Frank Marryat’s description of a voyage from Panama to San Francisco, 1851 [90]

It seemed that we had brought the yellow fever with us to Panama, or rather it appeared at the time of our arrival, and it was now spreading with great rapidity. Cholera also broke out, and deaths from one or the other of these causes became very numerous.

The people being panic-struck, a great rush was made for the Californian boats, of which there happened, at this time, to be very few.

So soon as I was able to move, there was but one small screw steamer in port, and as the place was daily becoming more unhealthy, I secured, by great favour, a cabin in her. Nothing could excuse the state in which this ship was put to sea, not even the panic; for she was not only ill-found in every respect, but was so crowded with passengers, that it was not until it was ascertained that there was scarcely standing-room for those on board that she tripped her anchor.

I had secured a dog-hole of a cabin, and was no sooner on board than my wife, worn out by fatigue and anxiety, was attacked by violent fever. There were two young doctors on board, but both were attacked shortly after we started. Then the epidemic (an aggravated intermittent fever) broke out among the passengers, who – crowded in the hold as thick as blacks in a slaver – gave way to fear, and could not be moved from the lower deck, and so lay weltering in their filth.

#### Sheldon F. Dudley’s Report of 1918 pandemic influenza on board Royal Navy vessels, 1921 [96]

The density of susceptible persons in a ship must be very great as compared with an assemblage of susceptible persons on shore. If, for example, we contrast the sleeping accommodation in a ship with that of a big institution, we find that in the ship hammock hooks are less than 2 ft. apart, whereas institution bed centres are rarely less than 8ft.apart. Even when head to toe slinging is insisted on in a ship the men’s heads must often be within 3ft. of each other. Now the volume of spray from a mouth at 3ft. is nearly twenty times that at 8ft. Therefore how much more readily will the man sleeping in a battleship’s mess-deck get a requisite dose of infectious material than the man sleeping in an institution ashore? Again we also hear of many men in a ship not ill enough to go off duty, and, thus being immobilized, wandering about among their fellows.

When we consider these points, and at the same time realize that a modern battleship, with its tiers of lumbered decks, its cramped accommodation, and its crew of often over 1,000 men, covers an area of less than one-fiftieth of a square mile, 1 do not think it possible to doubt the infective material must become so dense and diffused as to saturate the ship. That is to say, everyone on board receives a dose of the specific agent sufficient to cause influenza, unless he happens to be highly immune at that time to the strain of organism responsible for the epidemic. In a ship, the density of susceptible persons, the mass of infection, and the local migration are all so great that any diminution in one or more of these factors that may be produced by the use of sprays and gargles, by early isolation of cases and disinfection, is scarcely likely to diminish the rate of spread in a ship, once influenza has obtained a footing on board. And I think naval experience, where all these preventive measures have been vigorously employed, justifies this pessimism, as I have been unable to learn of any definite cases in which they did any good. In ships the outbreaks lasted ten days to three weeks; ashore the wave took about three months to pass over a locality. The longer wave period ashore was probably due to the lesser density of susceptible persons and infective sources, more time being required for the infection to hunt out all the susceptible individuals within its reach.

#### Text S3. Model Description and Equations

To achieve a more realistic depiction of the time course of infection, we make dwell times in state *E* and state *I* Erlang-distributed using the Linear Chain Trick [85]. This technique gives a unimodal distribution with a long right-hand tail, such that disease progression is relatively constrained in most individuals, but with occasional individuals experiencing substantially longer periods of incubation or infectiousness [86].

Individuals progress through *k*_*E*_ exposed states, *E*_1_, *E*_2_, …, *E*_*kE*_, and through *k*_*I*_ infectious states, *I*_1_, *I*_2_, …., *I*_*kI*_. We use *k*_*E*_ = *k*_*I*_ = 3 for all simulations.

The rate of progression from state *E*_*e*_ to state *E*_*e*+1_ and from state *E*_*kE*_ to state *I*_1_ is 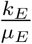, where *µ*_*E*_ represents the mean length of time that an individual spends in all exposed states. Similarly, the rate of progression from state *I*_*i*_ to state *I*_*i*+1_ and from state *I*_*kI*_ to state *R* is is 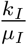, where *µ*_*I*_ represents the mean length of time that an individual spends in all infectious states.

We track infection across *g >* 1 transmission generations, where *E*_*n,e*_ and *I*_*n,i*_ denote *n*^th^-generation individuals in states *E*_*e*_ and *I*_*i*_ respectively. The 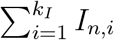 infectious individuals from generation *n* produce new exposed individuals *E*_*n*+1,1_, which represent the (*n* + 1)^st^ generation of infections.

The number of first-generation individuals, *n* = 1, is fixed at *t* = 0. For all simulations, we assume a single first-generation individual. We randomly assign this person to a state from *E*_1,1_, *E*_1,2_, …*E*_1,*kE*_ at the time of departure, *t* = 0.

To account for uncertainty and variation in the density dependence of shipboard contact rates, our model does not assume either classical density dependence or classical frequency dependence. Instead, we model the shipboard transmission with the equations:

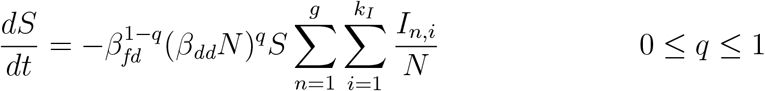

Here, *q* represents the degree of density dependence on board the ship, with *q* = 0 representing classical frequency dependence, *q* = 1 representing classical density dependence, and 0 *< q <* 1 representing intermediate modes of transmission. The parameters *β*_*dd*_ and *β*_*fd*_ modulate the intensity of transmission under each density dependence pole — intuitively, the proportion (*β*_*dd*_) and the raw number (*β*_*fd*_) of people on board ship that a single infected individual will infect per day, on average, in a fully-susceptible population. Since susceptible people can be infected by infectious people in any generation and at any stage of infection,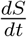 is proportional to the total number of infected people across all *g* transmission generations and all *k*_*I*_ infectious states.

The following equations show the deterministic analogue of our model. We implement continuous stochastic simulations in R using the Gillespie Algorithm, using the package GillespieSSA [84].

### Model Equations

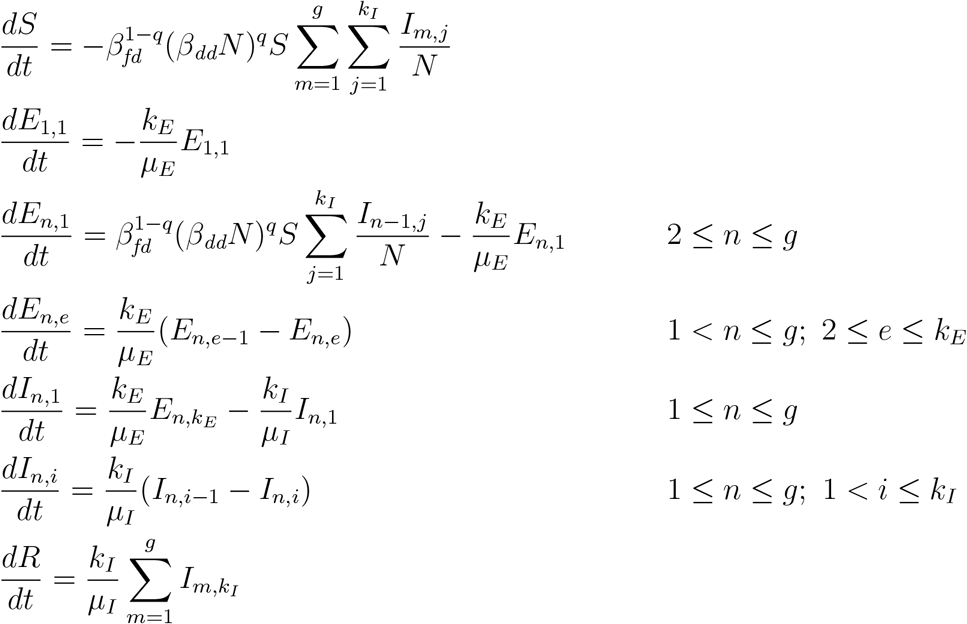

## Supplementary Figures

**Fig. S1.**
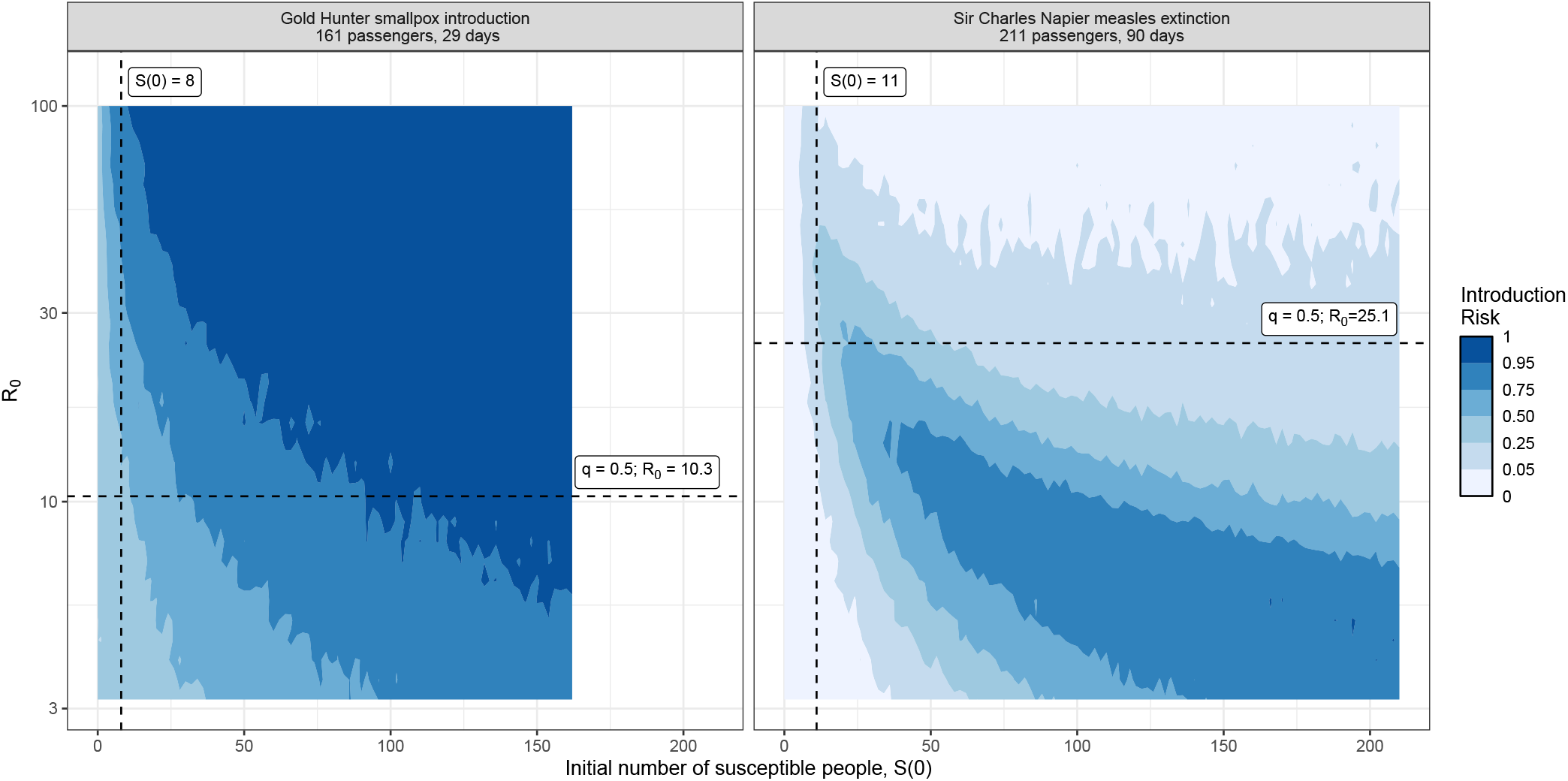
Introduction risk for documented pathogen outbreaks on board the *Gold Hunter* (smallpox) and the *Sir Charles Napier* (measles) by initial number of susceptibles, S(0) and by transmission rate, R_0_. All analyses assume a single initial index case (*e*_0_ = 1). X-axis limits reflect each ship’s population: 163 for the *Gold Hunter* and 211 for the *Sir Charles Napier* (Table S4). We use limits of [0,162] and [0,210] for the *Gold Hunter* and the *Sir Charles Napier*, respectively, where 0 represents no susceptible people other than the index case. Introduction probability represents the likelihood of sustained outbreaks across a journey of 29 days (*Gold Hunter*) and 90 days (*Sir Charles Napier*). Horizontal and vertical lines reflect, respectively, the values of *S*(0) and *R*_0_ used in Figure 4, Table 1, and Table 2.

**Fig. S2.**
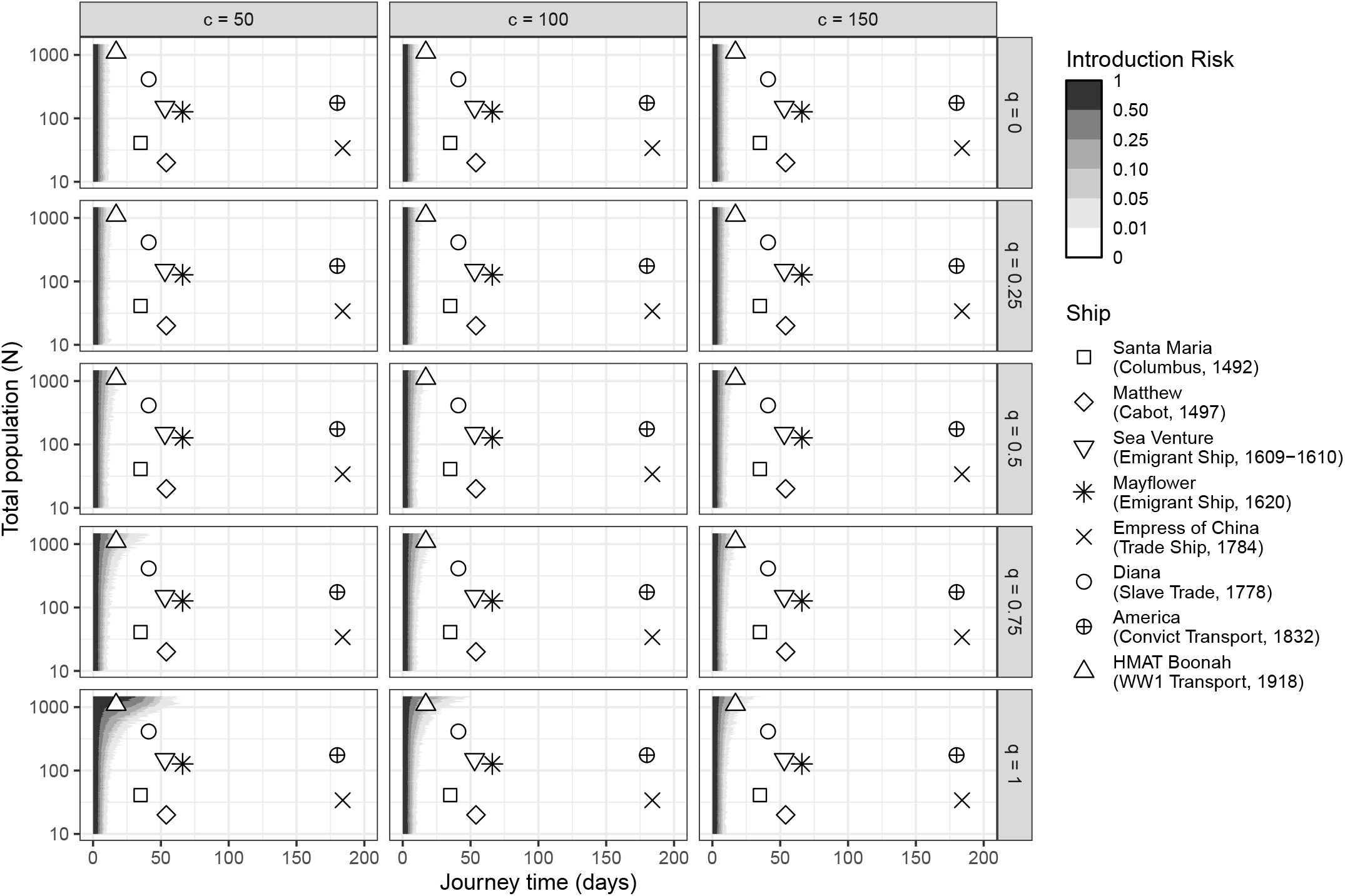
Sensitivity of influenza introduction risk to density dependence index, *q*, and to density dependence scaling constant, *c*. All analyses assume a single index case (*e*_0_ = 1), a population susceptibility rate *S*(0)*/N* = 0.05, and natural history parameters *µ*_*E*_, *µ*_*I*_ and *β*_*fd*_ as given in Table S1.

**Fig. S3.**
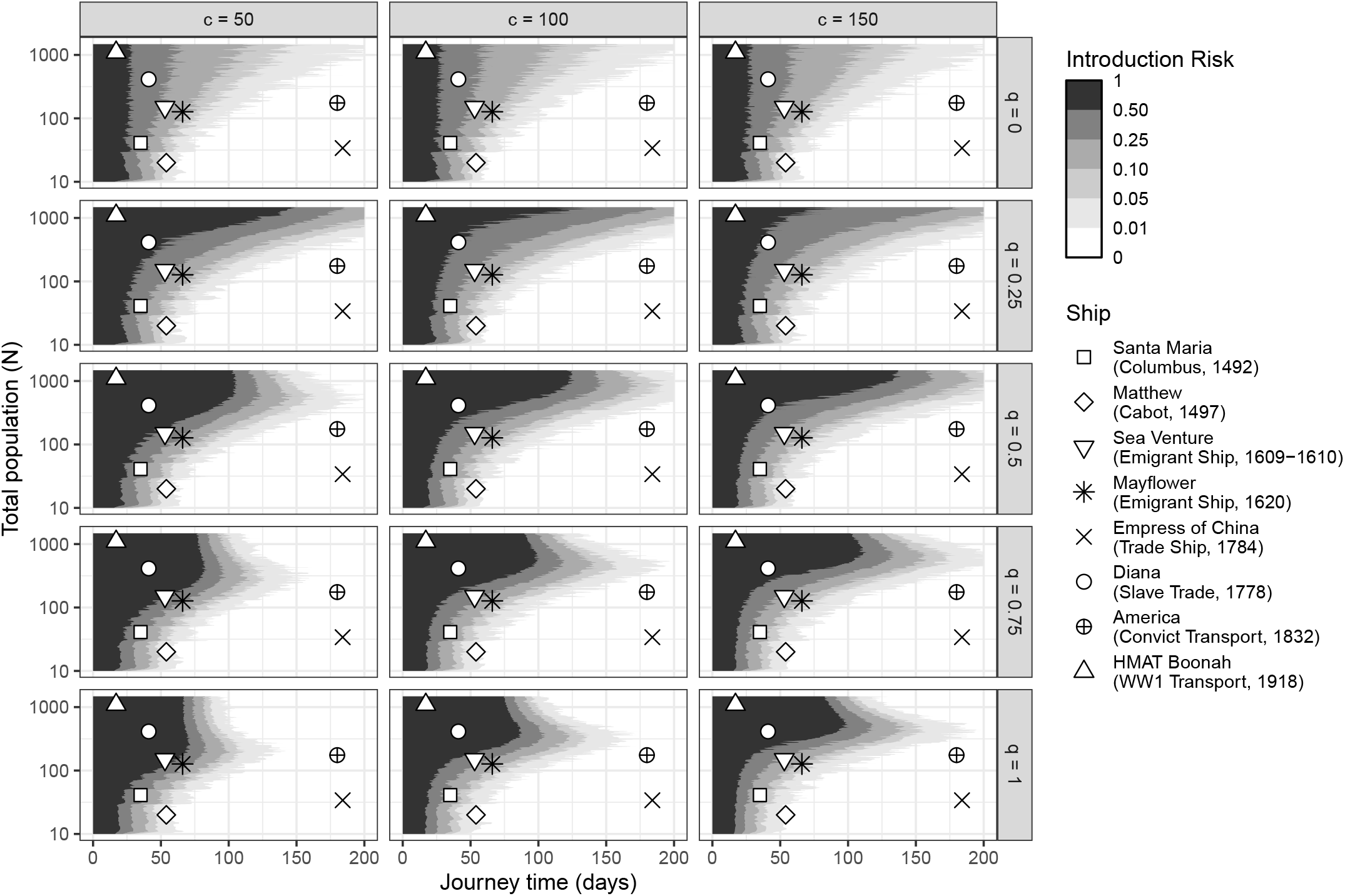
Sensitivity of measles introduction risk to density dependence index, *q*, and to density dependence scaling constant, *c*. All analyses assume a single index case (*e*_0_ = 1), a population susceptibility rate *S*(0)*/N* = 0.05, and natural history parameters *µ*_*E*_, *µ*_*I*_ and *β*_*fd*_ as given in Table S1.

**Fig. S4.**
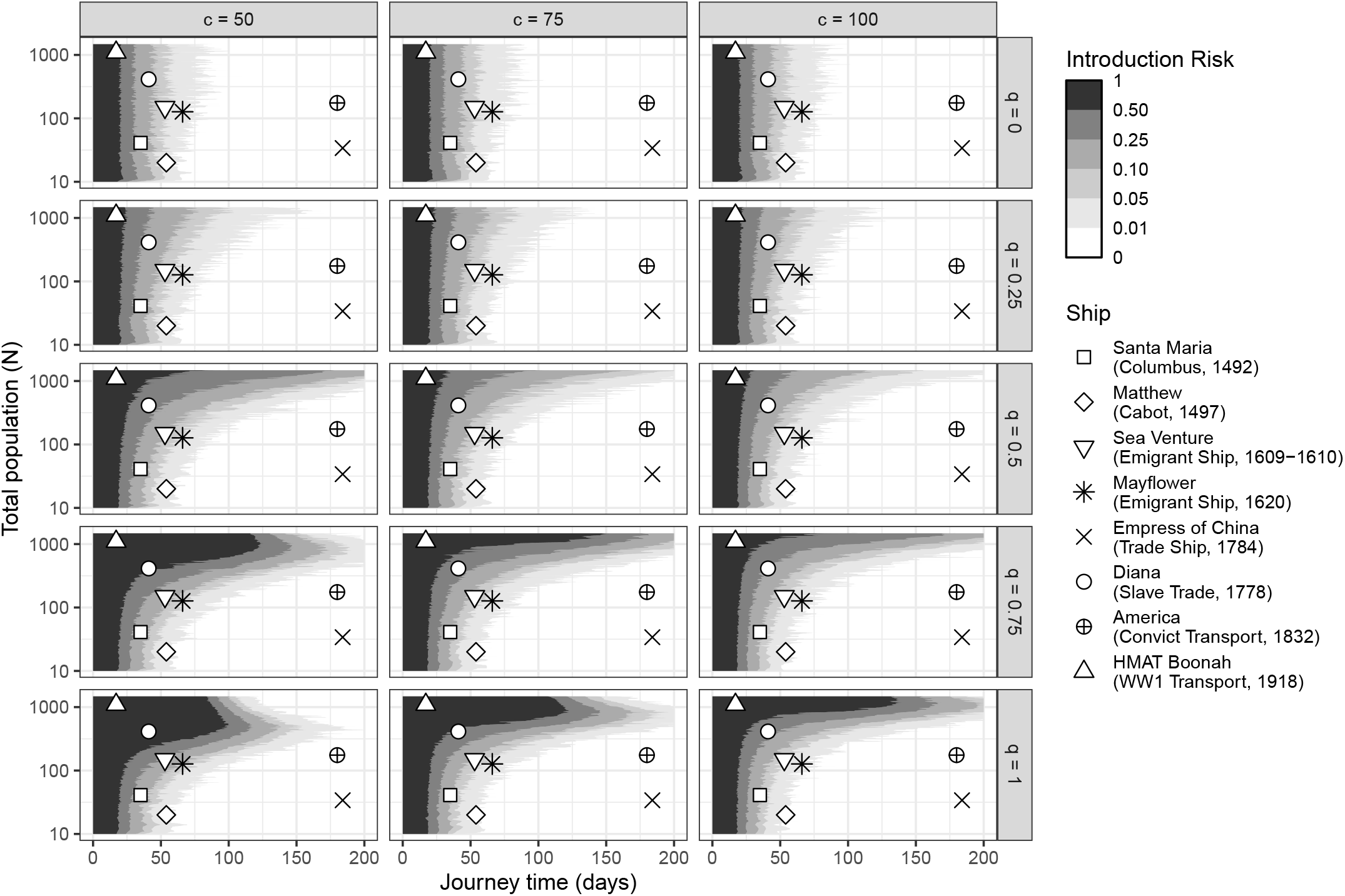
Sensitivity of smallpox introduction risk to density dependence index, *q*, and to density dependence scaling constant, *c*. All analyses assume a single index case (*e*_0_ = 1), a population susceptibility rate *S*(0)*/N* = 0.05, and natural history parameters *µ*_*E*_, *µ*_*I*_ and *β*_*fd*_ as given in Table S1.

**Fig. S5.**
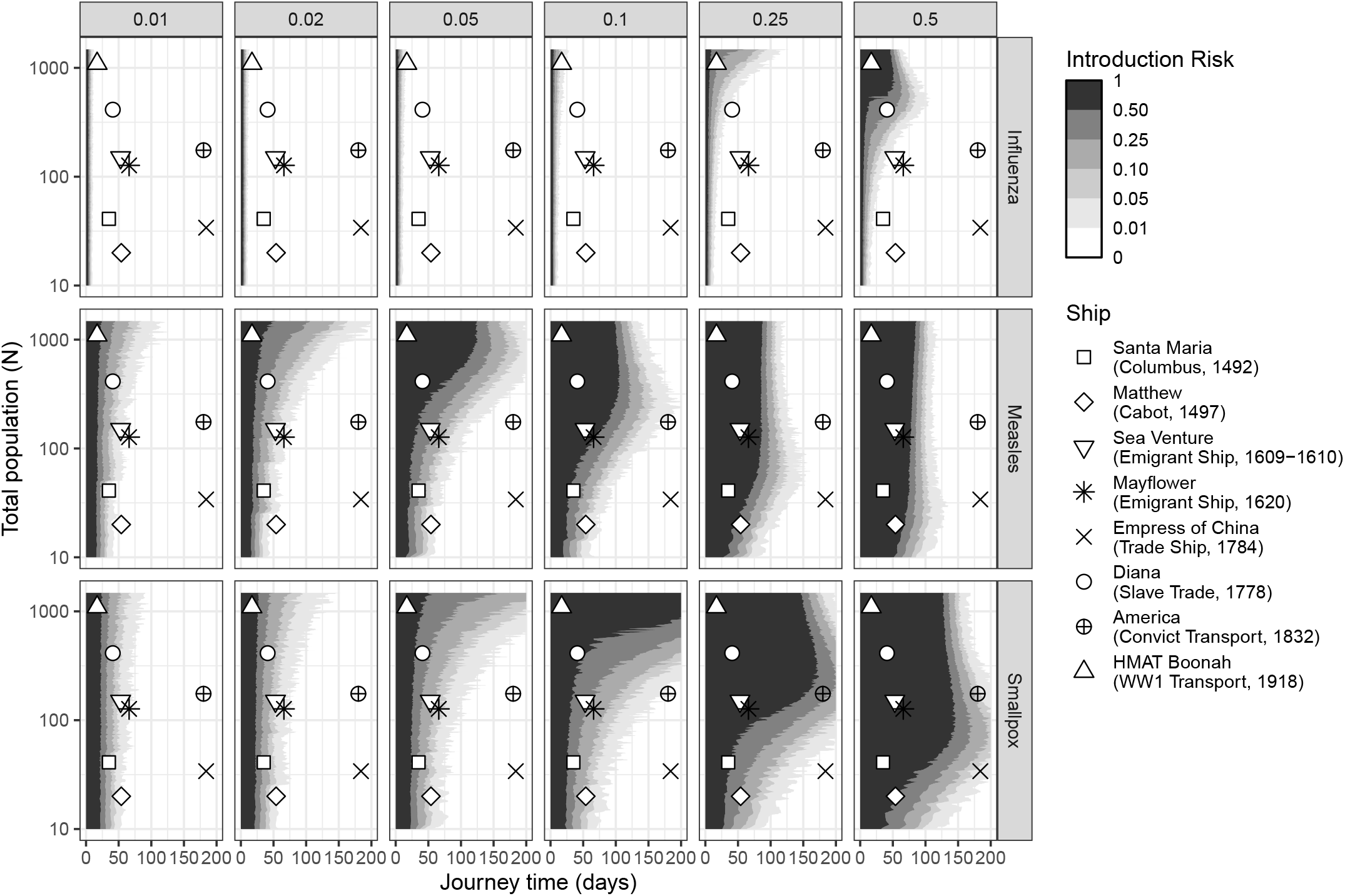
Sensitivity of introduction risk to initial population susceptibility, *S*(0)*/N*, by pathogen. All analyses assume a single index case (*e*_0_ = 1), intermediate density dependence (*q* = 0.5), a density dependence scaling constant *c* = 100, and pathogen-specific natural history parameters as given in Table S1.

**Table S1.** Natural History Parameters.

| Pathogen | Mean latent period (days) | Mean infectious period (days) | Typical Land $R_0$ | $\beta_{fd}$ used for Figure 4, Tables 1–2 | $\beta_{dd}$ used for Figure 4, Tables 1–2 | $R_0$ used for Figure 4, Tables 1–2 | Reference |
| --- | --- | --- | --- | --- | --- | --- | --- |
| Influenza | 2 | 3 | 1.5 | 0.5 | 0.005 | $0.05\sqrt{N}$ | [110–112] |
| Measles | 12 | 8 | 15 | 1.875 | 0.01875 | $0.1875\sqrt{N}$ | [110,113] |
| Smallpox | 12 | 17.5 | 7 | 0.4 | 0.004 | $0.04\sqrt{N}$ | [114–116] |

**Table S2.** San Francisco Port Arrivals Statistics, June 1850 – June 1852.

| Origin | Type | $n$ | Journey Time (days) | | | Number of Passengers | | |
| --- | --- | --- | --- | --- | --- | --- | --- | --- |
|  |  |  | Median | Range | Interdecile Range | Median | Range | Interdecile Range |
| Liverpool, UK | Sail | 18 | 189.5 | 150-300 | 162-245.5 | 4.5 | 1-18 | 2-18 |
| New York, USA* | Sail | 92 | 155.5 | 89-283 | 113.1-219.8 | 5 | 0-160 | 1.1-20.0 |
|  | Steam | 9 | 180 | 65-364 | 87.2-296 | 111 | 5-743 | 11.4-515.0 |
| Valparaíso, Chile | Sail | 100 | 57 | 23-190 | 45.0-74.2 | 7 | 0-175 | 1.0-39.0 |
| Panama** | Sail | 39 | 63 | 36-122 | 43.4-83.4 | 53 | 1-287 | 4.6-164.6 |
|  | Steam | 116 | 20 | 12-58 | 15.7-28.0 | 196 | 11-1050 | 50.6-527.4 |
| Oregon | Sail | 48 | 7 | 3-16 | 3.4-11.3 | 4 | 0-12 | 1.0-6.0 |
|  | Steam | 47 | 3 | 2-8 | 2.5-5.0 | 28 | 3-157 | 12.0-60.0 |
| Hawai'i | Sail | 76 | 22.5 | 14-80 | 16.0-31.5 | 5 | 0-142 | 2.0-22.0 |
|  | Steam | 1 | 13 | – | – | 18 | – | – |
| Sydney, Australia | Sail | 20 | 91 | 67-188 | 70.0-111.2 | 15.5 | 2-142 | 3.9-57.0 |
| Hong Kong | Sail | 14 | 58.5 | 33-95 | 46.7-84.0 | 201 | 1-553 | 14.5-378.4 |

\* For reasons that are unclear, four steam voyages originating in New York City had transit times longer than 200 days: the *SS New Orleans* (210 days), the *SS Sea Bird* (240 days), the *SS Goliah* (279 days) and the *SS Chesapeake* (364 days). The *SS Sea Bird* ran aground on San Martine on its way to California and had to be repaired [42]. Meanwhile, the extraordinarily long voyage of the *SS Chesapeake* astonished contemporary observers, with companies who had shipped goods on the vessel suing for damages on its arrival [117].
\*\* Voyages from Panama predate the Panama Canal, which opened in 1914. Prior to its construction, travellers from Europe and the east coast took a ship to the eastern coast of Panama, crossed the Panama Isthmus by land, and took a second ship from the west coast of Panama to San Francisco [118].

**Table S3.** Selected Historical Voyages, 1492-1918.

| Year | Vessel | Type | Purpose | Journey | Duration | Population | Reference |
| --- | --- | --- | --- | --- | --- | --- | --- |
| 1492 | *Santa María*<br>[Columbus's Second Voyage] | Sail | Exploration | San Sebastian de la Gomera, Canary Islands to San Salvador, present-day Bahamas | 35 days | 41-60 crew (estimates vary; we use the more recent estimate of 41) | [119] |
| 1497-98 | *Matthew*<br>[Cabot's Second Voyage] | Sail | Exploration | Bristol, England to a disputed location in northeastern North America, possibly present-day Maine, Newfoundland, or Nova Scotia | 54 days | 20 crew | [120] |
| 1609 | *Sea Venture* | Sail | Emigration | Bristol, England to present-day Bermuda | 53 days | 150 passengers and crew | [121] |
| 1620 | *Mayflower* | Sail | Emigration | Leiden, Holland to near Cape Cod, Massachusetts | 66 days | 102 passengers, 20-30 crew (we assume 25 crew) | [122] |
| 1784 | *Empress of China* | Sail | Trade | New York City to Macao | 184 days | 34 crew | [114] |
| 1778 | *Diana* | Sail | Slave ship | Iles de Los, present-day Guinea to Curaçao | 41 days | 413 enslaved people, 30 crew | [123] |
| 1832 | *America* | Sail | Convict transport | Unknown (likely United Kingdom to Australia) | 180 days | 175 crew and convicts | [36] |
| 1918 | HMAT *Boonah* | Steam | Troop ship | Durban, South Africa to Fremantle, Australia | 17 days | 164 crew, 931 troops | [124] |

**Table S4.** Known outbreaks of influenza, smallpox, and measles 1850–1918.

| Year | Vessel | Type | Pathogen | Journey | Journey Duration | N | Outbreak Duration | Cases | Reference |
| --- | --- | --- | --- | --- | --- | --- | --- | --- | --- |
| 1851 | *Gold Hunter* | Steam | Smallpox | Panama – San Francisco | 29 days | 163 | 29+ days | 1 case | [50] |
| 1852 | *Sir Charles Napier* | Sail | “Measles, dysentery, and fever” | Panama – San Francisco | 90 days | 211 | “about three weeks” (newspaper article); 54 days (final death) | 36 deaths (total for all outbreaks; no cause of death data available) | [49] |

